# Profiling serum oxylipin metabolites across melanoma subtypes and immunotherapy responders

**DOI:** 10.1101/2025.02.05.636637

**Authors:** Alexander C. Goodman, Kylie M. Michel, Morgan L. MacBeth, Jaqueline A. Turner, Richard P. Tobin, William A. Robinson, Kasey L. Couts

## Abstract

**Objectives:** This study investigates the relationship between serum oxylipin profiles and response to immune checkpoint inhibitor therapy in melanoma subtypes to identify potential metabolic biomarkers for treatment response.

**Methods:** In a retrospective cohort study, serum samples from 43 stage III and stage IV melanoma patients treated at the University of Colorado Hospital from 2010 to 2023 were analyzed via ultra-high-pressure liquid chromatography-mass spectrometry. Melanoma patients were treated anti-PD-1 monotherapy or combination immune checkpoint inhibitor therapy and response was assessed using RECIST 1.1 criteria.

**Results:** Using mass spectroscopy, we determined global oxylipin metabolite profiles are largely uniform pre-and post-treatment across melanoma subtypes including cutaneous, acral, mucosal, and uveal melanoma. Across subtypes, 33 oxylipin metabolites were analyzed, with limited variation observed overall. Prostaglandin J_2_ was more abundant in rare melanoma subtypes including acral, mucosal, and uveal melanoma compared to cutaneous melanoma.

**Conclusions:** Despite limited variation of serum oxylipin molecular species by subtype and response status, we observed significant differences in Prostaglandin J_2_ which could serve as a potential biomarker for immune checkpoint inhibitor therapy response in melanoma. However, further investigation is warranted to explore the role of oxylipins in immune response modulation.

## Introduction

Oxylipins are a family of fatty acids that have undergone at least one oxidation reaction and are typically derived from polyunsaturated fatty acids (PUFAs) like omega-3 and omega-6 fatty acids. These molecules are produced through three primary pathways: cyclooxygenases (COXs), lipoxygenases (LOXs), and cytochrome P450 enzymes (CYPs) [1]. Common omega-3 PUFAs, such as eicosapentaenoic acid (EPA) and docosahexaenoic acid (DHA), give rise to derivatives like hydroxyeicosapentaenoic acids (HEPEs), hydroxydocosahexaenoic acids (HDoHEs), and resolvin D1 (RvD1), which are pro-resolving and anti-inflammatory lipid mediators [2]. In contrast, omega-6 PUFAs, such as arachidonic acid (AA), produce metabolites like prostaglandins (PGs), thromboxanes (TXs), leukotrienes (LTs), and hydroxyeicosatetraenoic acids (HETEs), which are pro-inflammatory lipids [2]. Additional PUFAs, including dihomo-gamma-linolenic acid (DGLA) and linolenic acid (LA), also produce oxylipin derivatives that display both pro-and anti-inflammatory activities, further complicating the role of these molecules in inflammatory processes [1].

Previous reports show oxylipin molecular species, specifically prostaglandin E_2_ (PGE_2_), are elevated in melanoma. PGE_2_ is generated from COX-2, which has elevated expression in melanoma and is associated with increased proliferation, enhanced invasion, and a worse overall prognosis [3,4]. Additionally, the suppression of COX-2 leads to enhanced infiltration of CD8⁺ T cells and dendritic cells into the tumor microenvironment, suggesting COX-2 modulates adaptive immune responses [5]. PGE_2_ can significantly influence the efficacy of immune checkpoint inhibitor (ICI) therapy in cancers including melanoma. Elevated tumoral PGE_2_ levels are associated with immunotherapy resistance [6–8], and inhibition of COX rescue anti-PD-1 efficacy in resistant tumors [6]. Combination anti-CTLA-4 antibodies and prostaglandin E_2_ receptor 4 (EP4) antagonists, which block PGE_2_ signaling, enhances tumor cytoreduction [7]. Even omega-3 fatty acids enhance ICI response, whereas omega-6 fatty acids impede efficacy. Higher plasma levels of long-chain fatty acids, especially omega-3s, are associated with better ICI responses in melanoma patients [9], suggesting that circulating lipid profiles may influence immunotherapy efficacy and serve as potential biomarkers for predicting patient response and potentially reprogram inflammatory and adaptive immune responses during ICI therapy.

While prostaglandins have been widely studied in melanoma, the contributions of other oxylipins, including leukotrienes and LOX-derived metabolites, remain less understood [4]. *In vitro* melanoma treatment with LOX inhibition, specifically 15-LOX inhibitors, results in reduced proliferation, decreased cell viability, and enhanced cytotoxicity compared to COX inhibitors [10]. Interestingly, melanoma-reprogrammed Schwann cells have been found to upregulate 12-LOX, 15-LOX, and COX-2, leading to increased production of pro-tumorigenic oxylipins including PGE₂ and lipoxins A₄/B₄, and subsequent anti-tumor T-cell activation [11]. These findings suggest that both COX and LOX pathways are instrumental in melanoma progression and immune evasion. Rarer subtypes of melanoma include acral melanoma (AM), occurring on the palms, soles, fingers, toes, and nail units; mucosal melanoma (MM), which arises from mucosal surfaces; and uveal melanoma (UM), originating from melanocytes in the iris, ciliary body, and choroid of the eye [12]. Existing studies indicate that these rare melanomas exhibit lower response rates to ICI therapy. For instance, patients with acral melanoma show an objective response rate of 32% to combination ICI therapy, which is lower than that observed in cutaneous melanoma [13]. Mucosal melanoma responds less effectively to both single-agent and combination ICI therapy, with a progression-free survival of 5.9 months compared to 11.7 months in cutaneous melanoma patients receiving combination therapy [14]. Uveal melanoma patients demonstrate an even lower objective response rate of 18% to combination ICI therapy [13]. While genomic and transcriptomic differences among these melanoma subtypes have been identified [15–17], research has predominantly focused on these genetic variations, with little attention given to metabolomics, particularly lipidomics. Notably, a study by Vilbert and colleagues (2023) demonstrated that cutaneous melanoma (CM), MM, and UM exhibit distinct metabolic profiles, specifically in lipid composition. They analyzed choline-containing phospholipids and sphingolipids, finding that serum levels of these lipids were lower in patients with MM and higher in those with CM and UM [12]. Similarly, de Bruyn et al. (2023) found significant differences in the metabolome of UM patients compared to healthy controls, particularly in metabolites associated with malignant processes [18]. These studies suggest that metabolic differences, especially in lipid profiles, exist among melanoma subtypes and may contribute to their distinct biological behaviors.

Given the diverse responses to ICI therapy among melanoma subtypes and the potential influence of oxylipins on therapeutic outcomes, there is a critical need to understand how circulating oxylipin profiles correlate with ICI responses across different melanoma types. Peripheral blood analysis offers a less invasive and more accessible means to identify biomarkers predictive of treatment response, circumventing the challenges associated with tumor biopsies. This study investigates the association between circulating levels of oxylipins and the response to immune checkpoint blockade in patients with different melanoma subtypes. By profiling oxylipins in peripheral blood, we seek to identify metabolic signatures that correlate with ICI therapy outcomes. Specifically, differences in pro-inflammatory and anti-inflammatory oxylipins may provide insights into the mechanisms of ICI resistance and predict therapeutic efficacy as potential biomarkers for personalized treatment strategies.

## Methods

### Study Population

This is an exploratory retrospective observational cohort study of patients with advanced melanoma treated with ICI therapy at the University of Colorado Hospital. Serum samples were collected between 2010 and 2023. Samples were collected in red-top tubes directly in the hospital, processed immediately, and stored in a-80C freezer until use. For the purposes of our study, we focused on the PD-1 immune checkpoint blockers. Patients were included in the study if they received either pembrolizumab (Keytruda) or nivolumab (Opdivo) monotherapy or combination therapy [nivolumab (Opdivo) + ipilimumab (Yervoy)]. When possible, we collected a pre-treatment ICI therapy serum sample and an on-treatment ICI therapy sample at the time of their first follow up. If serum samples were drawn more than 6 months prior to treatment start, they were excluded from treatment-related analysis. The serum samples were obtained as part of the International Melanoma Biorepository and Research Laboratory at the University of Colorado Cancer Center. Patients were consented under the approval of the Colorado Institutional Review Board (IRB# 05-0309). These patient studies were conducted according to the Declaration of Helsinki, Belmont Report, and U.S. Common Rule.

Response was defined according to RECIST 1.1 criteria [19]. We classified a patient as a responder if they showed a complete response (CR) or partial response (PR) and non-responders were designated as stable disease (SD) or progressive disease (PD) at time of follow-up. Follow-up time average around one year and ranged from zero to six years.

### Oxylipin Quantification by UHPLC-MS

Oxylipin profiles were measured in melanoma human serum samples using mass spectrometry. Serum samples were diluted 1:10 with cold 5:3:2 MeOH:ACN:H2O (v/v/v) containing deuterated standards (Cayman Chemical) each at a final concentration of 6 ng/uL ( Supplementary Table 1, Supplementary Digital Content 1). Samples were vortexed vigorously for 30 minutes at 4°C, then centrifuged for 10 minutes at 18,213 RCF. Using 10 uL injection volumes, the supernatants were analyzed by ultra-high-pressure liquid chromatography coupled to mass spectrometry (UHPLC-MS). Metabolites were resolved across a 1.7 um, 2.1 x 100 mm Waters Acquity BEH column using a 7-minute gradient previously described [20].

Following data acquisition,.raw files were converted to.mzXML using the RawConverter application. Metabolites were annotated and peaks integrated based on intact mass, 13C isotope pattern, and retention times in conjunction with the deuterated standards and KEGG database. Peaks were integrated using El-Maven (Elucidata). Quality control was assessed as using a technical mix of all samples run at beginning, end, and middle of each sequence as previously described [21].

### Experimental Data and Statistical Analysis

Absolute quantification of study samples was reported. All studied samples were normalized to internal standards. All statistical analyses and visualizations were conducted in R (R version 4.4.0, http://www.r-project.org). The significance value was set to alpha = 0.05.

Summaries for demographics are presented as N (%) for categorical variables and mean (SD) and range for continuous variables. Before any analysis was conducted, testing for a normal distribution of the data was conducted using the Shapiro-Wilk test of normality. Given most measured study samples were not normally distributed in all comparisons (Supplementary Digital Content 2), we elected to use non-parametric testing.

## Results

### Patient Clinical Characteristics

Forty-three melanoma patients with at least stage III disease were included within the study cohort (Table 1) and represented the four major subtypes of melanoma including cutaneous (*n* = 6), acral (*n* = 7), mucosal (*n* = 23), and uveal (*n* = 7). Mucosal melanoma was further subdivided into three anatomic locations, anorectal (*n* = 8), nasopharyngeal (*n* = 8), and vulvovaginal (*n* = 7). A total of 26 female and 17 male melanoma patients ranging from 39 to 92 years old with an average age of 63 years old represented the entire cohort. The frequency of patients receiving anti-PD-1 immunotherapy or combination therapy (anti-PD-1 and anti-CTLA4) ranged from 14.3% to 62.5% across melanoma subtypes. Rates of ICI responders ranged from 0% (AM) to 66.7% (CM).

**Table 1.**
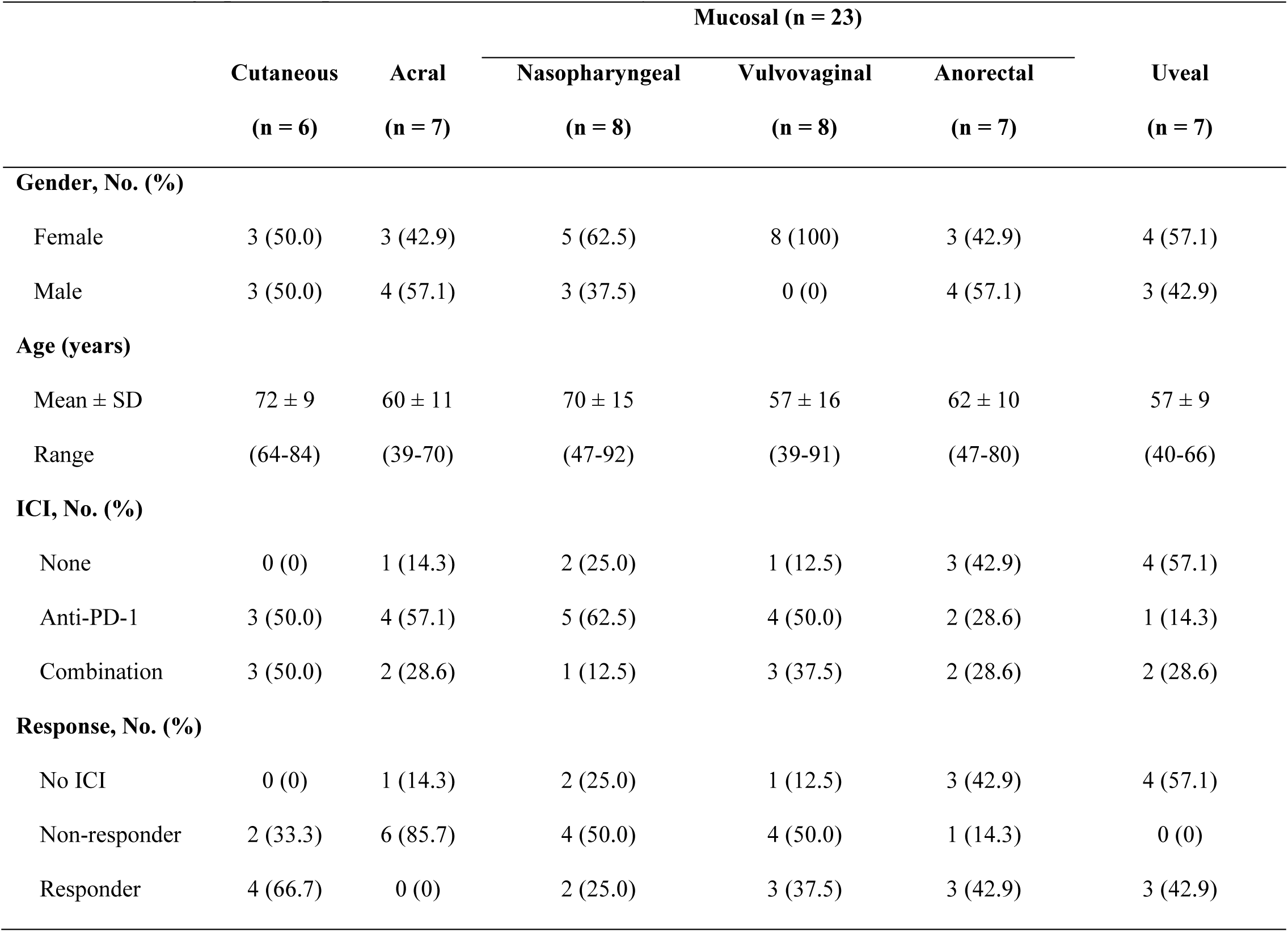
Demographics of patients included in the study.

### Mass spectrometry evaluation of oxylipin levels in patient serum

A total of 33 oxylipin metabolites were analyzed using mass spectrometry (Supplementary Table 1, Supplementary Digital Content 1). Each of the lipid metabolites were further classified into six polyunsaturated fatty acids (PUFAs), 15 arachidonic acid (AA) derivates, four docosahexaenoic acid (DHA) derivatives, one eicosapentaenoic acid (EPA) derivative, and seven linoleic acid (LA) derivatives (Supplementary Figure 1, Supplementary Digital Content 1). We performed an unsupervised principal component analysis (PCA) for all samples and identified 4 outlier samples which were removed from further analysis (Supplementary Figure 2a, Supplementary Digital Content 1). A heatmap generated from unsupervised clustering of all specimens showed polyunsaturated fatty acids clustered separately from other fatty acids as expected (Supplementary Figure 2b, Supplementary Digital Content 1).

### Baseline serum oxylipins across melanoma subtypes

To evaluate which oxylipins differed between the melanoma subtypes at baseline, oxylipin concentrations were analyzed in ICI therapy naïve patient serum samples. First, we performed an unsupervised PCA analysis. The first two principal components, PC1 and PC2, accounted for greater than 45% of the variation in the data and demonstrated a significant amount of overlap melanoma subtypes (Figure 1a). We calculated the loading scores for the first and second principal components to determine which lipids contributed to the most variation. Assessment of the top 10 oxylipins with highest loading scores showed that a significant portion of these oxylipins we part of LA metabolism (Figure 1b-c). Unsupervised hierarchical clustering revealed no distinct global clustering pattern based on melanoma subtype (Figure 1d) which was further supported by post-hoc analysis using Kruskal-Wallis rank sum test with Dunn’s test (Supplementary Digital Content 3).

**Figure 1.**
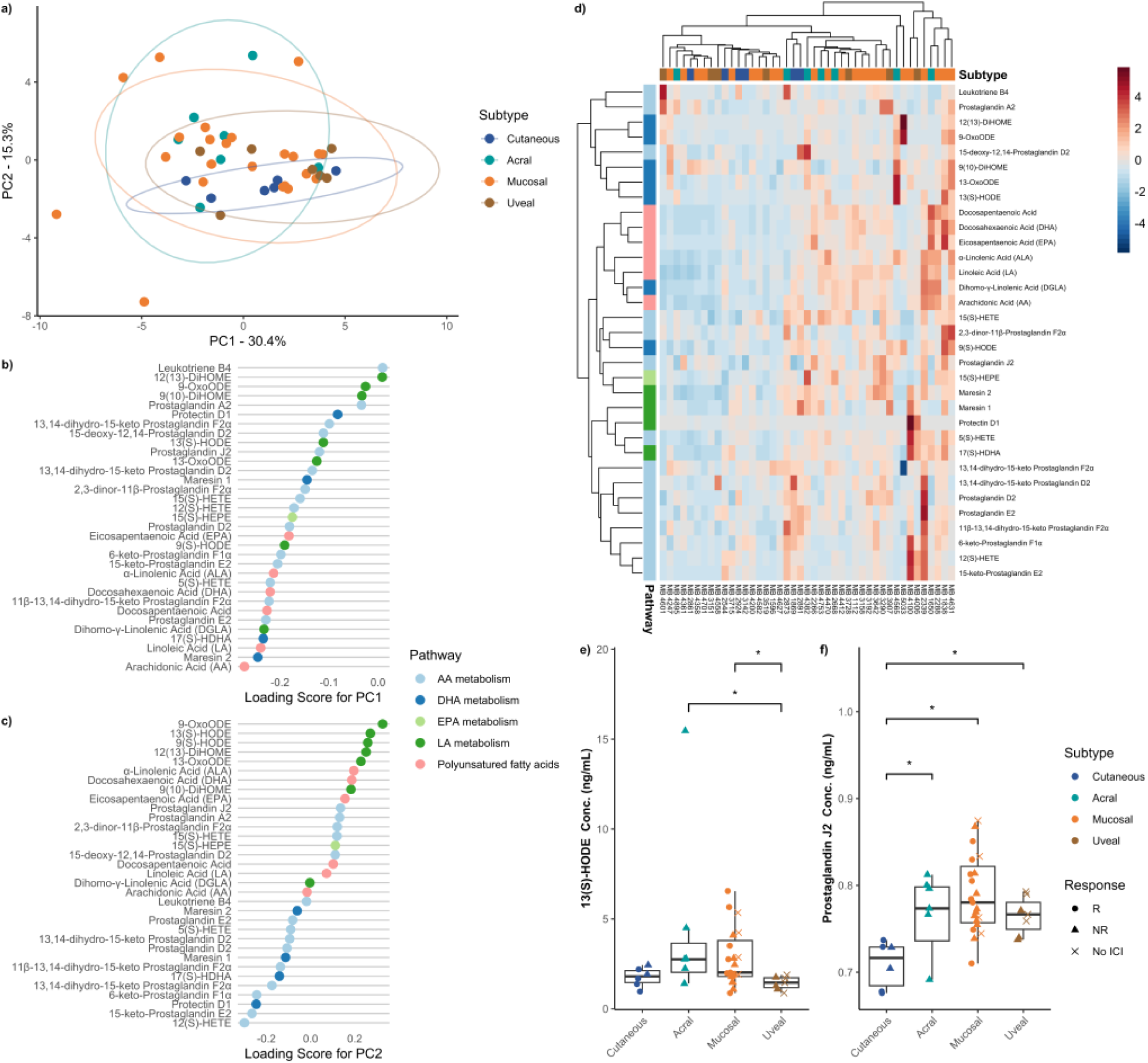
Summary of comparison between melanoma subtypes. (a) Principal component analysis discriminating against healthy donors and melanoma subtypes. The explained variance of each component is indicated on the corresponding axis. Plot of loading scores for (b) PC1 and (c) PC2 of the principal component analysis. Plotted values are absolute to signify magnitude of effect each oxylipin has on the principal component. (d) Heatmap demonstrating differences in serum oxylipins concentration between melanoma subtypes. Boxplot of absolute serum concentration of (e) 13(S)-HODE and (f) prostaglandin J_2_ for all melanoma subtypes prior to any ICI therapy. Significant differences are indicated by an asterisk (*p* ≤ 0.05). Trend differences are indicated by a cross (*p* ≤ 0.100).

Upon further stratifying the oxylipins, we did find 13(S)-HODE was enriched in acral and mucosal melanoma compared to uveal melanoma (*H* = 9.584, *p* = 0.022). Prostaglandin J_2_ (PGJ_2_) was significantly more abundant in rare melanomas, including acral, mucosal, and uveal as compared to cutaneous melanoma (*H* = 13.715, *p* = 0.003). Both acral (*p* = 0.028) and mucosal (*p* = 0.028) melanoma patients had significantly higher serum levels of 13(S)-HODE than uveal melanoma (Figure 1e). PGJ_2_ was higher in acral (*p* = 0.048), mucosal (*p* = 0.001), and uveal (*p* = 0.050) melanoma patients when compared to cutaneous (Figure 1f). There was a notable trend towards decreased 11β-13,14-dihydro-15-keto Prostaglandin F_2α_ in rare melanomas compared to cutaneous (*H* = 6.378, *p* = 0.095) and enrichment in 13-OxoODE in acral and mucosal melanoma (*H* = 6.667, *p* = 0.083) (Supplementary Figure 3, Supplementary Digital Content 1). While not reaching significance, cutaneous (*p* = 0.131), acral (*p* = 0.129), and mucosal (*p* = 0.175) melanoma patients had higher levels of serum 11β-13,14-dihydro-15-keto Prostaglandin F_2α_ at baseline than uveal melanoma patients. Similarly, there were no significant differences in serum 13-OxoODE across the melanoma subtypes, but acral (*p* = 0.111) and mucosal (*p* = 0.111) patients showed higher levels than uveal patients. Together, these results suggest that while there are global profile differences in oxylipin metabolism across subtypes, AA and LA metabolite derivatives and 13(S)-HODE and PGJ_2_ are differentially enriched in rare melanoma subtypes.

### Baseline serum oxylipins across mucosal melanoma anatomic locations

Next, we questioned if circulating oxylipin abundancy differed across mucosal melanoma anatomic locations. ICI therapy naïve samples from patients with anorectal, nasopharyngeal, and vulvovaginal mucosal melanoma were compared and two-dimensional PCA accounted for greater than 53% of the variation in the data with clustered sample overlap despite different mucosal etiologies (Figure 2a). Assessment of the top 10 oxylipins with highest loading scores showed that a significant portion of these oxylipins we part of the PUFAs (Figure 2b-c). Hierarchical clustering analysis did not delineate distinct metabolic profiles despite different anatomic locations (Figure 2d). Further analysis on individual metabolites recapitulated an enriched abundancy of PGJ_2_ in mucosal melanomas compared to cutaneous but no significant difference across mucosal anatomic location (*H* = 14.854, *p* = 0.002) (Figure 2e). No other individual metabolites showed any notable enrichments (Supplementary Digital Content 4; Supplementary Figure 4, Supplementary Digital Content 1). Cutaneous melanoma patients showed significantly lower baseline serum levels of PGJ_2_ than vulvovaginal (*p* = 0.025) and nasopharyngeal (*p* = 0.025) mucosal melanoma patients, with anorectal (*p* = 0.001) mucosal melanoma patients having the highest levels and most statistically different from cutaneous (Figure 2e). Thus, oxylipins metabolism shows variation in the AA metabolic pathway with increased abundancy of PGJ_2_ across all mucosal melanoma anatomic primary sites.

**Figure 2.**
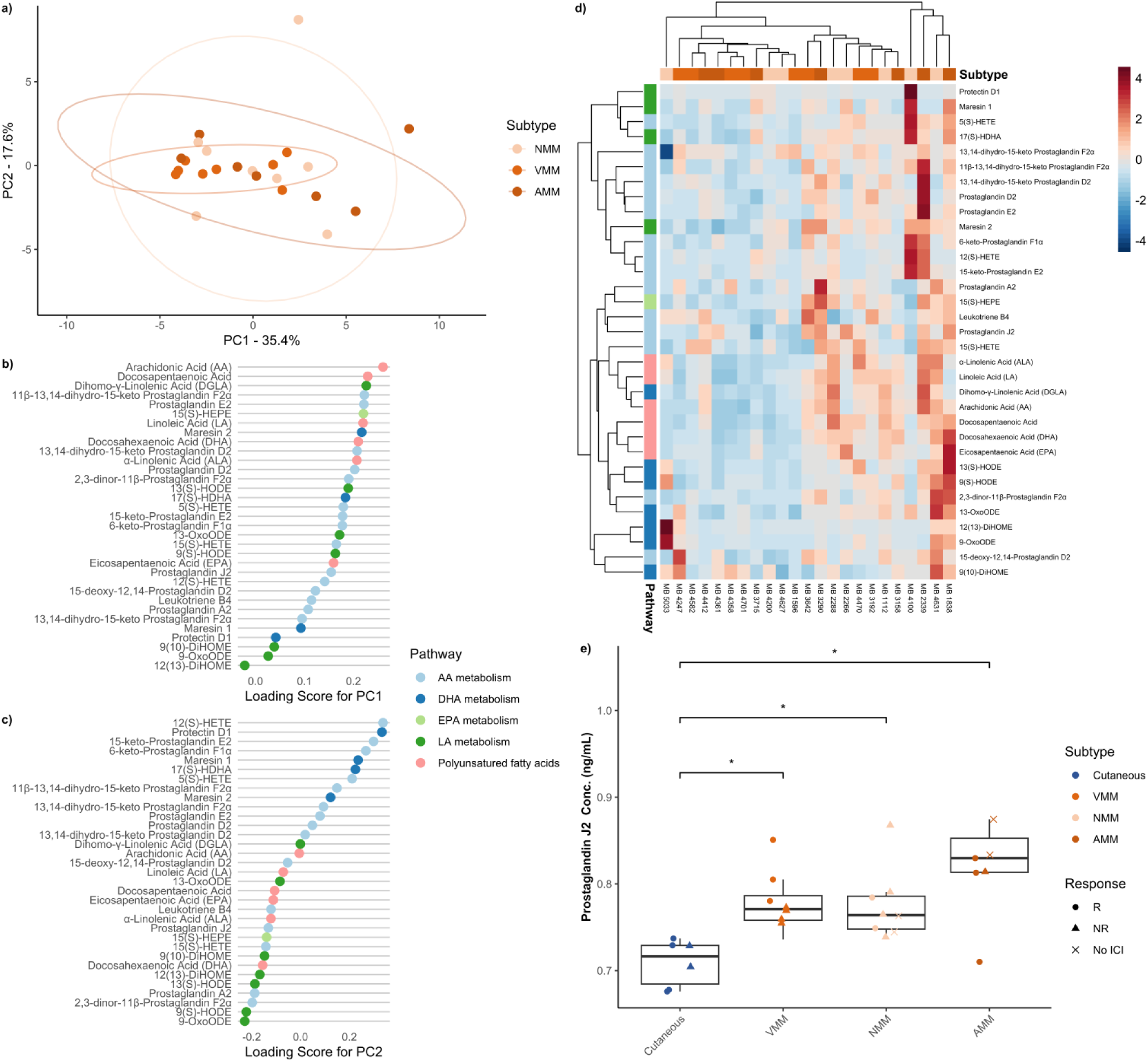
Summary of comparison between mucosal melanoma anatomic locations. **(a)** Principal component analysis discriminating against mucosal melanoma locations. The explained variance of each component is indicated on the corresponding axis. Plot of loading scores for **(b)** PC1 and **(c)** PC2 of the principal component analysis. Plotted values are absolute to signify magnitude of effect each oxylipin has on the principal component. **(d)** Heatmap demonstrating differences in serum oxylipins concentration between mucosal melanoma locations. **(e)** Boxplot of absolute serum concentration of prostaglandin J_2_ for cutaneous melanoma and mucosal melanoma locations prior to any ICI therapy. Significant differences are indicated by an asterisk (*p* ≤ 0.05). Trend differences are indicated by a cross (*p* ≤ 0.100).

### Baseline serum oxylipins between immune checkpoint therapy responders and non-responders across melanoma subtypes

Next, we looked at the differences in baseline, pre-treatment serum oxylipins between ICI therapy responders (R) and non-responders (NR) across the four melanoma subtypes. Serum samples were divided up by subtype and ICI response and compared using PCA. Samples from patients that did not go on to receive ICI therapy were excluded in the analysis. Dimension reduction analysis demonstrated overlap of individual patient samples and overall uniform distribution of the data variation despite subtype and response status where two-dimensional PCA accounted for greater than 46% of the variation (Figure 3a). Assessment of the top 10 oxylipins with highest loading scores, showed components were comprised of AA, DHA, an LA metabolites (Figure 3b-c). Unsupervised hierarchical clustering did not distinguish response status based on global metabolic profile (Figure 3d), however, some mucosal samples clustered based on enrichment of PUFAs. Individual metabolites analysis showed a significantly increased abundancy of PGJ_2_ in both R and NR rare melanomas (*H* = 12.066, *p* = 0.034). Specifically, cutaneous melanoma patients that responded to immune checkpoint therapy showed a trend toward lower levels of baseline serum PGJ_2_ than mucosal responders (p = 0.065) (Figure 3e). While not statistically significant, 2,3-dinor-11β-Prostaglandin F_2α_ was trending to be higher in cutaneous NR patient serum samples (*H* = 9.699, *p* = 0.084). There were no significant or trend differences for 2,3-dinor-11β-Prostaglandin F2α (Figure 3f) in the post-hoc analysis. No other individual metabolites showed any notable enrichments (Supplementary Digital Content 5; Supplementary Figure 5, Supplementary Digital Content 1). These findings support the previous findings that while there are no significant global metabolic profile changes, individual metabolites distinguish rare melanoma subtypes compared to cutaneous melanomas. Despite that PGJ_2_ is enriched in rare melanomas, it does discriminate responder status.

**Figure 3.**
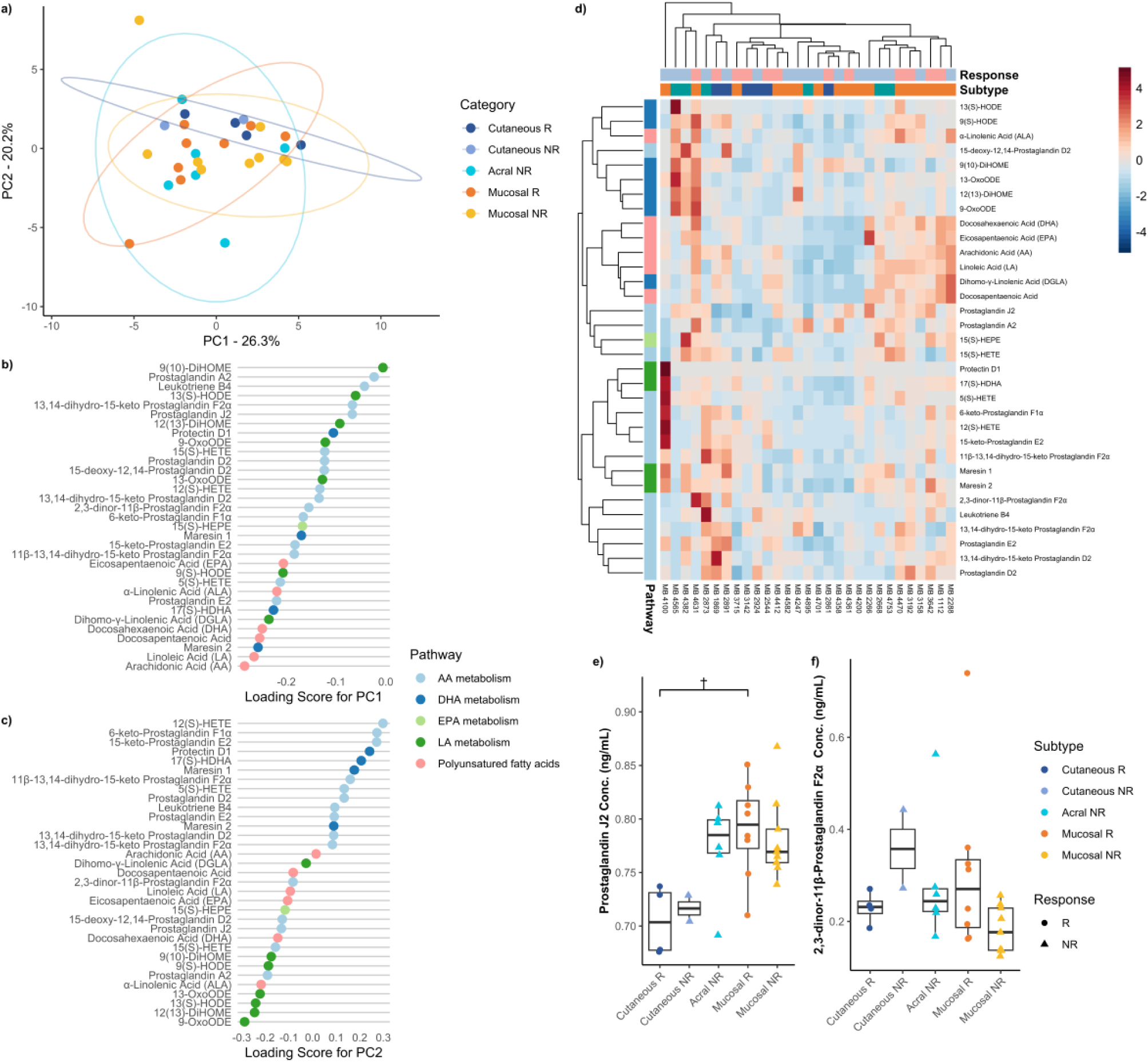
Summary of comparison between ICI therapy responders and non-responders for each melanoma subtype. **(a)** Principal component analysis discriminating against ICI therapy responders and non-responders across melanoma subtypes. The explained variance of each component is indicated on the corresponding axis. Plot of loading scores for **(b)** PC1 and **(c)** PC2 of the principal component analysis. Plotted values are absolute to signify magnitude of effect each oxylipin has on the principal component. **(d)** Heatmap demonstrating differences in serum oxylipins concentration between ICI therapy responders and non-responders across melanoma subtypes. Boxplot of absolute serum concentration of **(e)** prostaglandin J_2_ and **(f)** 2,3-dinor-11β- Prostaglandin F_2α_ for ICI therapy responders and non-responders across melanoma subtypes prior to any ICI therapy. Acral responders were not included in these plots as there were no acral responders in our study population. Significant differences are indicated by an asterisk (*p* ≤ 0.05). Trend differences are indicated by a cross (*p* ≤ 0.100).

### Difference between pre-treatment and on-/post-treatment oxylipins across melanoma subtypes

Lastly, we questioned how serum oxylipin levels change from pre-treatment ICI therapy vs on-treatment or post-treatment ICI therapy across the four melanoma subtypes. Uveal melanoma subtypes were excluded in this analysis as since on-/post-treatment samples were not available. Additionally, acral responders were not included in this analysis there was only one matched sample for this group. Pathway hierarchical clustering demonstrated clustering according to oxylipin pathway, particularly for on AA, LA, PUFA, and DHA metabolites (Figure 4a). However, no consistent patterns were discerned between pre-and on/post-treatment samples according to sample or response.

**Figure 4.**
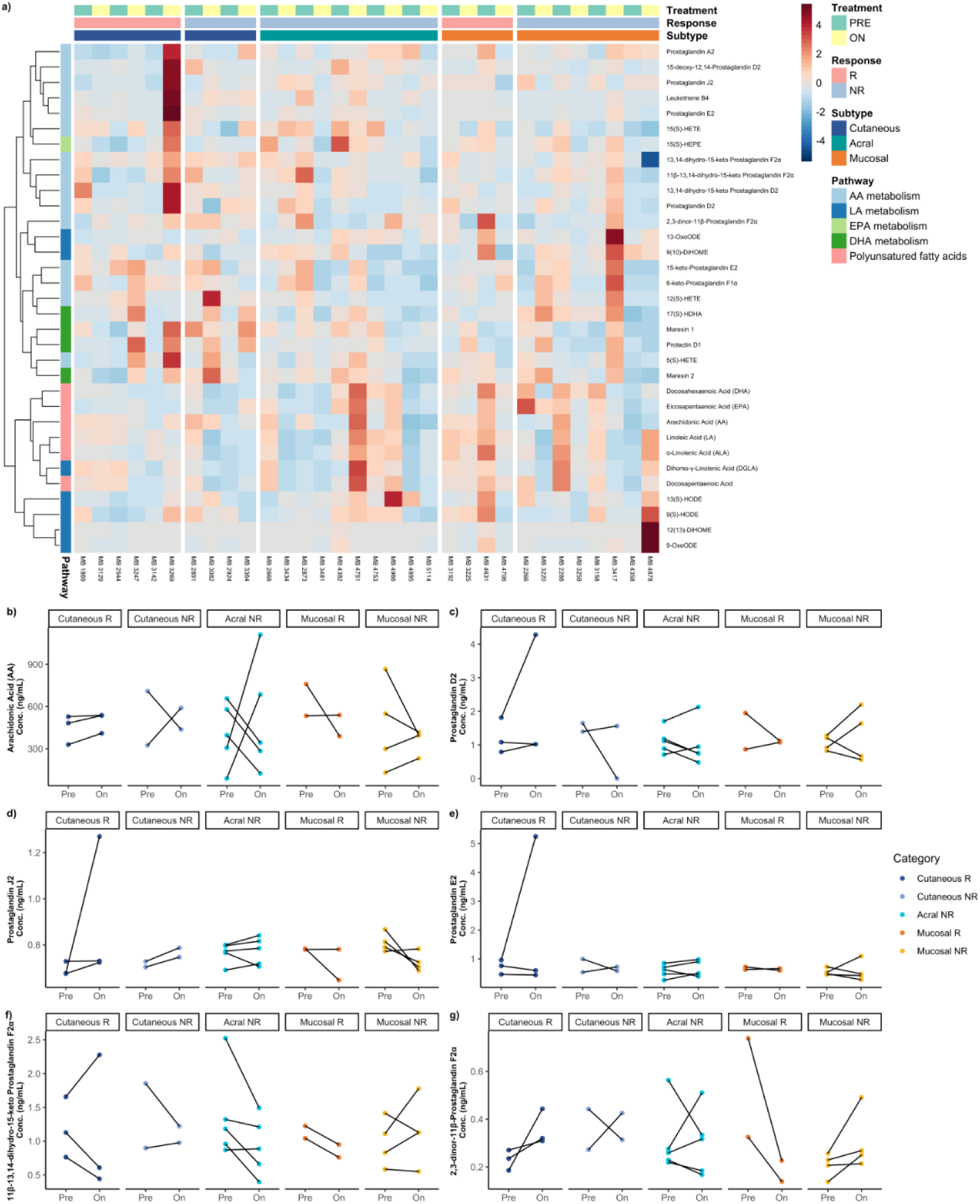
Summary of comparison between matched pre-and on/post-treatment samples in ICI therapy responders and non-responders for each melanoma subtype. **(a)** Heatmap demonstrating differences in matched pre-and on/post-treatment serum oxylipins concentration between ICI therapy responders and non-responders across melanoma subtypes. Line graphs of arachidonic acid-related metabolism oxylipins for matched pre-and on/post-treatment samples absolute serum concentration of **(b)** arachidonic acid, **(c)** prostaglandin D_2_, **(d)** prostaglandin J_2_, **(e)** prostaglandin E2, **(f)** 11β-13,14-dihydro-15-keto Prostaglandin F_2α_, and **(g)** 2,3-dinor-11β-Prostaglandin F_2α_ for ICI therapy responders and non-responders across melanoma subtypes prior to any ICI therapy. The “On” column was comprised of on-treatment and post-treatment samples.

We used a paired Wilcoxon signed-rank test to test for a difference in pre-treatment and on-/post-treatment serum samples across responders and non-responders for all melanoma types (Supplementary Digital Content 6). Given AA and LA metabolism drove a most of the variability, we assessed cancer-associated oxylipins in the AA pathway (Figure b-f) and the LA pathway (Supplementary Figure 6, Supplementary Digital Content 1). We did not find any significant or trend differences between pre-treatment and on-/post-treatment serum samples across all the measured oxylipins for any of the melanoma types.

## Discussion

While ICI therapy has revolutionized the way we treat melanoma and other cancers, a large portion of patients fail to respond to immunotherapy. Here, we investigated whether oxylipin profiles in serum samples of melanoma patients for the major melanoma subtypes correlated with response to ICI therapy. We observed limited variation in global oxylipin metabolite profiles across the melanoma subtypes and ICI therapy response. However, our data does suggest individual metabolites are enriched in rare melanoma subtypes. Further the minimal variation that was observed could represent differences in polyunsaturated fatty acids or arachidonic acid metabolites. Particularly, we found that ICI response is most influenced by polyunsaturated fatty acids. Interestingly, we repeatedly found differences in AA metabolism specifically between the cutaneous and the rarer melanoma subtype in that patients with cutaneous melanoma tended to have higher serum levels of these metabolites than patients with mucosal melanoma. However, given the little variation between our samples, more studies are needed to determine the how influential these highlighted differences truly are.

We identified one oxylipin, Prostaglandin J_2_ (PGJ_2_), that recurrently showed statistical differences both across melanoma subtypes and ICI responses. We found that this oxylipin is lower in cutaneous melanoma patients than acral, mucosal, and uveal melanoma patients. When we looked at PGJ_2_ in mucosal anatomic locations, we found that this oxylipin was higher in vulvovaginal, nasopharyngeal, and anorectal than cutaneous melanoma. Lastly, we found that CM responders had lower baseline levels of this oxylipin than mucosal responders.

PGJ_2_ and its derivative, 15-deoxy-Δ12,14-prostaglandin J_2_ (15d-PGJ_2_), have shown promise in cancer research, particularly for melanoma. Studies reveal that PGJ_2_ can inhibit cancer cell proliferation by inducing cell cycle arrest and apoptosis, primarily through activation of the peroxisome proliferator-activated receptor-gamma (PPARγ) pathway, which regulates genes that control cell growth and differentiation, making PGJ_2_ a candidate for targeting cancer cell survival mechanisms [22]. Additionally, PGJ_2_ has cytotoxic properties that enable it to disrupt proteins essential for tumor growth, which has been explored in melanoma as well as in breast, colon, and prostate cancers [23]. PGJ_2_ also plays a complex role in immunity and is capable of modulating the tumor immune microenvironment. Its anti-inflammatory properties can reduce chronic inflammation often linked to tumor progression, potentially inhibiting the supportive environment cancer cells need [24]. This anti-inflammatory effect also points to PGJ_2_’s potential in immunotherapy; by tempering inflammatory responses, PGJ_2_ might complement immune checkpoint inhibitors, such as anti-PD-1 and anti-CTLA-4 therapies, reducing side effects and possibly enhancing efficacy [25]. Additionally, PGJ_2_ may help activate immune cells, like macrophages, to adopt an anti-tumor role, suggesting it could “reprogram” the immune system to better target cancer [26]. Altogether, PGJ_2_’s roles in inhibiting tumor growth, modulating inflammation, and enhancing immune responses make it a promising candidate for both direct cancer therapy and as an adjunct in immunotherapy, especially for melanoma. However, further research is needed to fully realize its therapeutic potential and application in clinical settings.

The main limitation to our study was small sample size. We included rare melanoma patients for which sample collection is inherently less frequent and difficult to obtain large cohorts of specimens at a single institution. In the future, a multi-institutional approach would be beneficial for increasing sample numbers of rare melanoma subtype patients and ICI treated patients to increase statistical power for our initial observation in trending differences in polyunsatured fatty acids and metabolites in the arachidonic acid pathway.

## Conclusion

Immunotherapy has significantly improved clinical outcomes for patients with late-stage melanoma, yet a substantial portion of patients fail to respond to these treatments. The variability in responses to ICI therapy, both among individual patients and across different melanoma subtypes, underscores the need to explore the influence of circulating factors such as oxylipins on therapeutic outcomes. This study highlights the intricate role of oxylipin metabolism in melanoma and its potential to affect ICI efficacy. In particular, targeting specific metabolites like PGJ_2_ may open new avenues for personalized treatment strategies, especially for patients with rare melanoma subtypes. Further research is essential to deepen our understanding of these relationships and to translate these findings into improved clinical practice.

## Supporting information

Supplementary Digital Content 2

Supplementary Digital Content 3

Supplementary Digital Content 4

Supplementary Digital Content 5

Supplementary Digital Content 6

## Acknowledgements

We first thank the melanoma patients from the CU Cutaneous Oncology clinic for their consent and providing blood specimens used in this study. We are grateful to J. Haines and other members of the University of Colorado School of Medicine Metabolomics Core for conducting UHPLC-MS. We would like to thank all the members of the Center for Rare Melanomas for the helpful discussions. This study was supported by funding from the Tracks & Special Programs Department at Rocky Vista University to A.C.G. along with funds from the Moore Family Foundation and Patten-David Foundation.

## Author Contributions

A.C.G., K.L.C, W.A.R., and J.A.T conceived the study. A.C.G conducted statistical analyses and wrote the manuscript. K.M.M. assisted with patient sample preparation and M.L.M. assisted with statistical analysis design.

## List of Supplemental Digital Content

Supplemental Digital Content 1. PDF

Supplemental Digital Content 2. Xlsx

Supplemental Digital Content 3. Xlsx

Supplemental Digital Content 4. Xlsx

Supplemental Digital Content 5. Xlsx

Supplemental Digital Content 6. Xlsx

