## Supplementary Table and Figures for "Profiling serum oxylipin metabolites across melanoma subtypes and immunotherapy responders"

**Index**

Supplementary Figures and Tables Page

Supplementary Table 1 3
Supplementary Figure 1 4
Supplementary Figure 2 5
Supplementary Figure 3 6
Supplementary Figure 4 7
Supplementary Figure 5 8
Supplementary Figure 6 9

| Compound | Formula | Pathway | Parent | MedRt | Polarity | Standard |
|---|---|---|---|---|---|---|
| 11β-13,14-dihydro-15-keto Prostaglandin F2α | C20H34O5 | AA metabolism | 353.23 | 3.239 | [M-H]- | 11β-13,14-dihydro-15-keto Prostaglandin F2α d9 |
| 12(S)-HETE | C20H32O3 | AA metabolism | 319.23 | 4.477 | [M-H]- | 12(S)-HETE-d8 |
| 13,14-dihydro-15-keto Prostaglandin D2 | C20H32O5 | AA metabolism | 351.22 | 3.395 | [M-H]- | 13-14-dihydro-15-keto Prostaglandin D2-d9 |
| 13,14-dihydro-15-keto Prostaglandin F2α | C20H34O5 | AA metabolism | 353.23 | 3.297 | [M-H]- | 13,14-dihydro-15-keto Prostaglandin F2α/E1 d4 |
| 15-deoxy-12,14-Prostaglandin D2 | C20H30O4 | AA metabolism | 333.21 | 3.889 | [M-H]- | 13,14-dihydro-15-keto Prostaglandin D2 d4 |
| 15-keto-Prostaglandin E2 | C20H30O5 | AA metabolism | 349.20 | 3.054 | [M-H]- | 15-keto-Prostaglandin E2 d8 |
| 15(S)-HETE | C20H32O3 | AA metabolism | 319.23 | 4.331 | [M-H]- | 15(S)-HETE-d8 |
| 2,3-dinor-11β-Prostaglandin F2α | C18H30O5 | AA metabolism | 325.20 | 2.674 | [M-H]- | 2,3-dinor-11β-Prostaglandin F2α d4 |
| 5(S)-HETE | C20H32O3 | AA metabolism | 319.23 | 4.736 | [M-H]- | 5(S)-HETE-d8 |
| 6-keto-Prostaglandin F1α | C20H34O6 | AA metabolism | 369.23 | 2.588 | [M-H]- | 6-keto Prostaglandin F1a-d4 |
| Leukotriene B4 | C20H32O4 | AA metabolism | 335.22 | 3.726 | [M-H]- | Leukotriene B4-d4 |
| Prostaglandin A2 | C20H30O4 | AA metabolism | 333.21 | 3.731 | [M-H]- | Prostaglandin A2-d4 |
| Prostaglandin D2 | C20H32O5 | AA metabolism | 351.22 | 3.109 | [M-H]- | Prostaglandin D2-d9 |
| Prostaglandin E2 | C20H32O5 | AA metabolism | 351.22 | 3.012 | [M-H]- | Prostaglandin E2-d9 |
| Prostaglandin J2 | C20H30O4 | AA metabolism | 333.21 | 3.53 | [M-H]- | Prostaglandin J2/B2 d4 |
| 17(S)-HDHA | C22H32O3 | DHA metabolism | 343.23 | 4.352 | [M-H]- | 17(S)-HDHA-d5 |
| Maresin 1 | C22H32O4 | DHA metabolism | 359.22 | 3.794 | [M-H]- | Maresin 1 d5 |
| Maresin 2 | C22H32O4 | DHA metabolism | 359.22 | 3.945 | [M-H]- | Maresin 2 d5 |
| Protectin D1 | C22H32O4 | DHA metabolism | 359.22 | 3.702 | [M-H]- | Protectin D1 d5 |
| 15(S)-HEPE | C20H30O3 | EPA metabolism | 317.21 | 4.163 | [M-H]- | 15(S)-HEPE-d5 |
| 12(13)-DiHOME | C18H34O4 | LA metabolism | 313.24 | 4.013 | [M-H]- | 12(13)-DiHOME d4 |
| 13-OxoODE | C18H30O3 | LA metabolism | 293.21 | 4.31 | [M-H]- | 13-OxoODE-d3 |
| 13(S)-HODE | C18H32O3 | LA metabolism | 295.23 | 4.31 | [M-H]- | 13(S)-HODE-d4 |
| 9-OxoODE | C18H30O3 | LA metabolism | 293.21 | 4.386 | [M-H]- | 9-OxoODE-d3 |
| 9(10)-DiHOME | C18H34O4 | LA metabolism | 313.24 | 3.939 | [M-H]- | 9(10)-DiHOME d4 |
| 9(S)-HODE | C18H32O3 | LA metabolism | 295.23 | 4.379 | [M-H]- | 9(S)-HODE-d4 |
| Dihomo-γ-Linolenic Acid (DGLA) | C20H34O2 | LA metabolism | 305.25 | 5.732 | [M-H]- | Dihomo-y-Linolenic Acid-d6 |
| Arachidonic Acid (AA) | C20H32O2 | Polyunsatured fatty acids | 303.23 | 5.612 | [M-H]- | Arachidonic Acid-d8 |
| Docosahexaenoic Acid (DHA) | C22H32O2 | Polyunsatured fatty acids | 327.23 | 5.509 | [M-H]- | Docosahexaenoic Acid-d5 |
| Docosapentaenoic Acid | C22H34O2 | Polyunsatured fatty acids | 329.25 | 5.658 | [M-H]- | Docosapentaenoic Acid-d5 |
| Eicosapentaenoic Acid (EPA) | C20H30O2 | Polyunsatured fatty acids | 301.22 | 5.275 | [M-H]- | Eicosapentaenoic Acid-d5 |
| Linoleic Acid (LA) | C18H32O2 | Polyunsatured fatty acids | 279.23 | 5.658 | [M-H]- | Linoleic Acid-d11 |
| α-Linolenic Acid (ALA) | C18H30O2 | Polyunsatured fatty acids | 277.22 | 5.333 | [M-H]- | a-Linolenic Acid-d5 |

**Supplementary Table 1**. Characteristics and UHPLC-MS parameters of oxylipins.

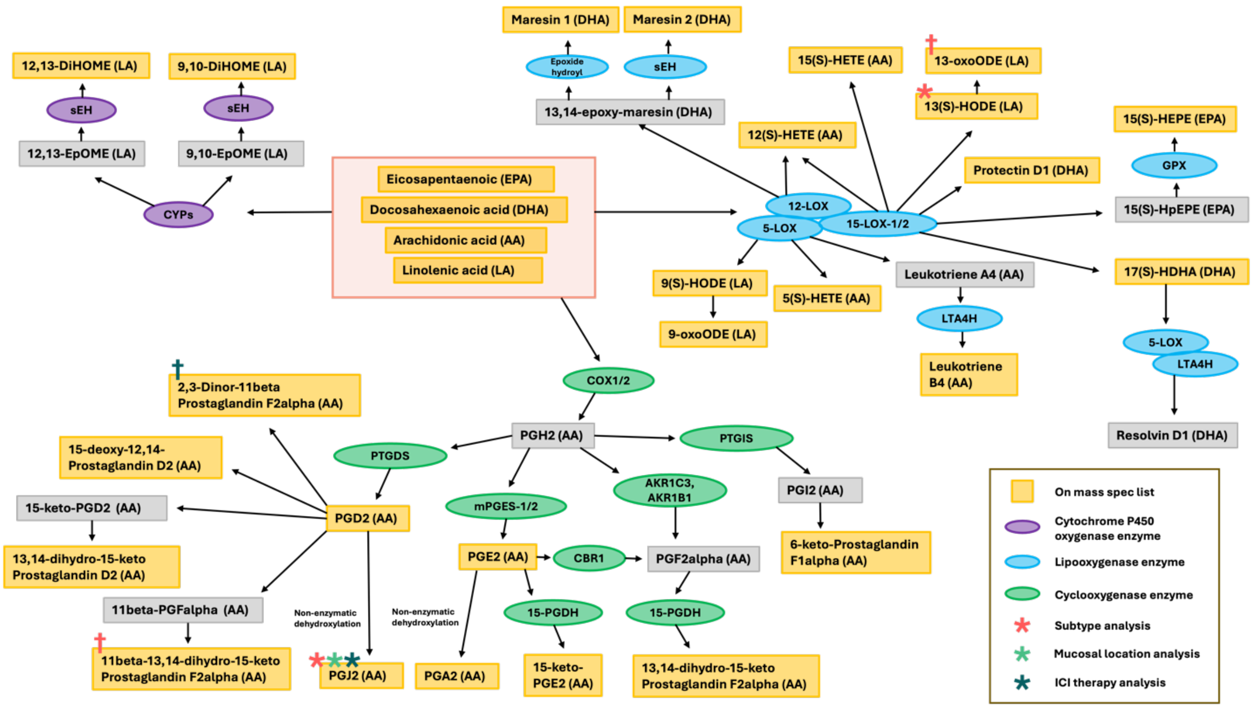


**Supplementary Figure 1.** Diagram of oxylipin classes analyzed in this study. Figure adapted and modified from Chistyakov and colleagues (2022). Asterisk above the listed oxylipin indicates a significant main affect in its respective analysis ($p < 0.05$). Cross above listed oxylipin indicates a trend effect in its respective analysis ($p < 0.100$).

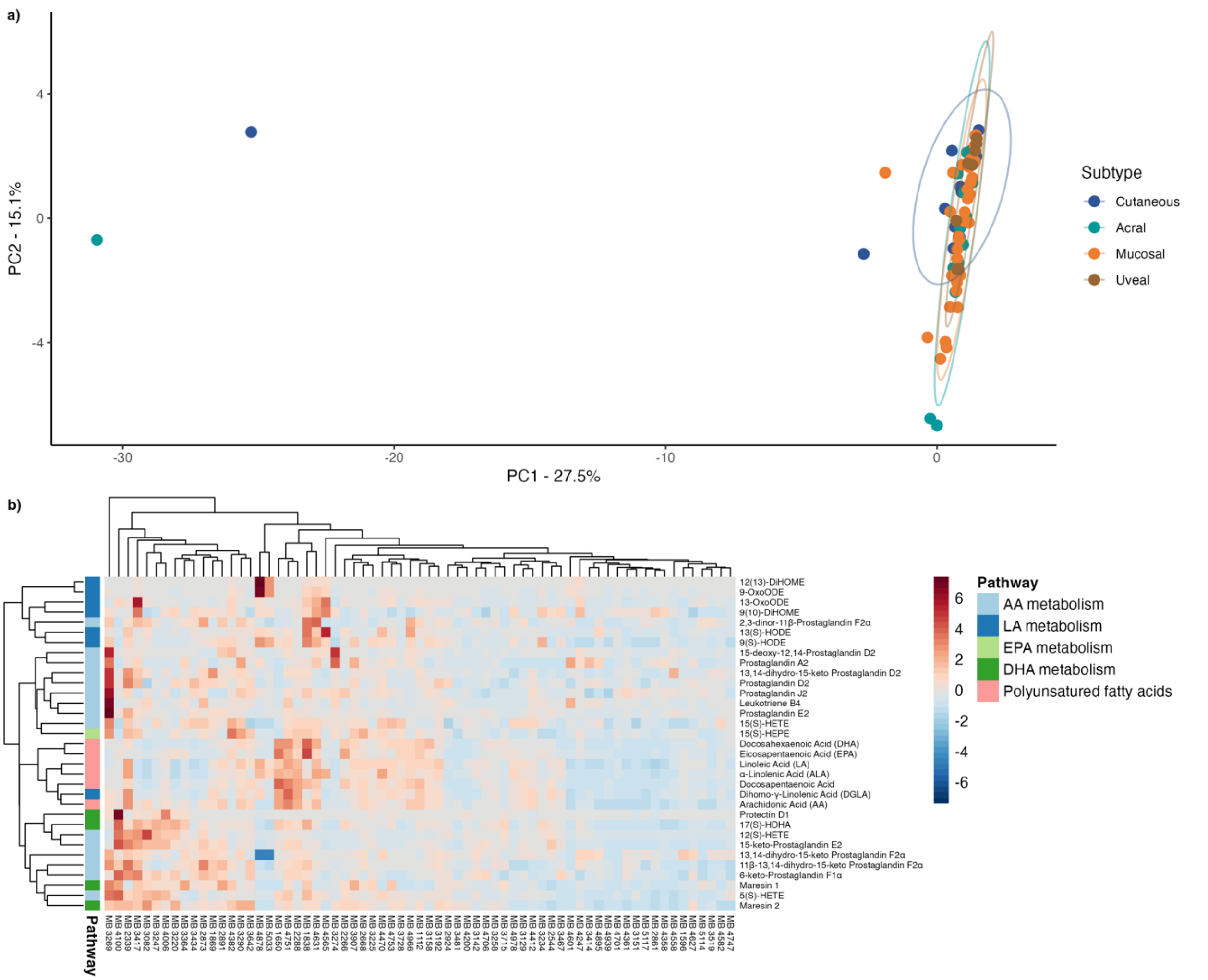


**Supplementary Figure 2.** Mass spectrometry screen for oxylipin levels in melanoma patient serum samples. **(a)** Principal component analysis of all samples that underwent mass spectrometry analysis. Plot demonstrates four outliers from total dataset, which were removed from further analysis. **(b)** Heatmap of all serum samples included in the study.

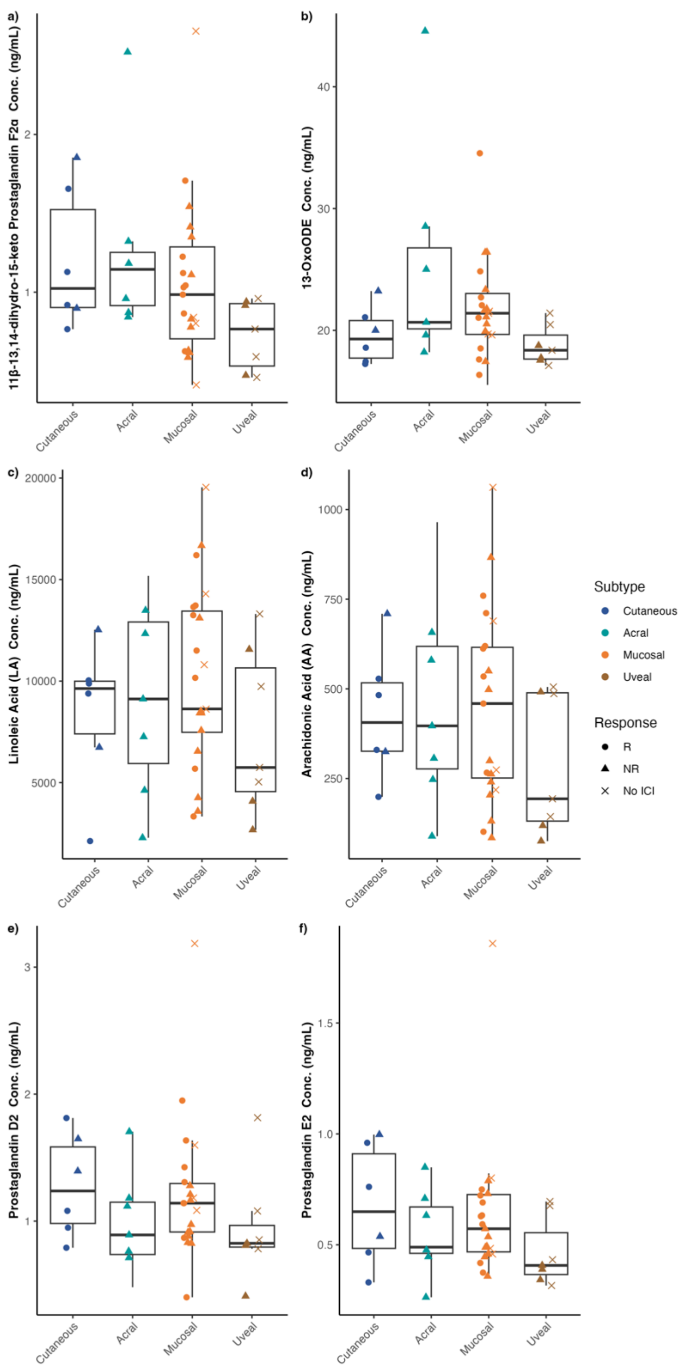


**Supplementary Figure 3.** Comparisons with trend effects of subtype and cancer-associated oxylipins for comparison. Boxplots of absolute serum concentration of **(a)** 11β-13,14-dihydro-15-keto Prostaglandin F2α, **(b)** 13-oxoODE, **(c)** linoleic acid, **(d)** arachidonic acid, **(e)** prostaglandin D2, and **(f)** prostaglandin E2 for all melanoma subtypes prior to any ICI therapy. Significant differences are indicated by an asterisk ($p \leq 0.05$). Trend differences are indicated by a cross ($p \leq 0.100$).

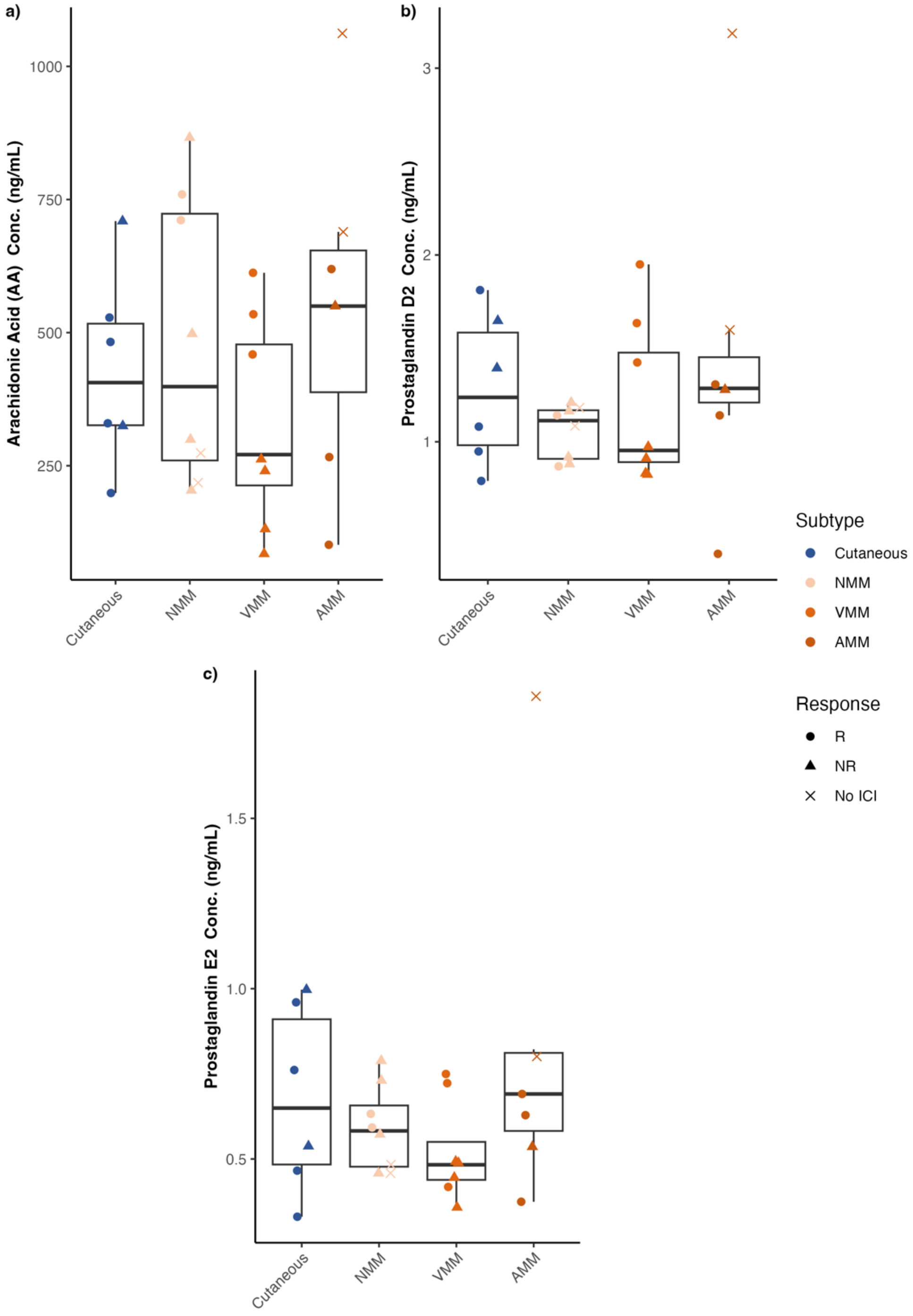


**Supplementary Figure 4.** Comparisons of cancer-associated oxylipins between mucosal melanoma anatomic locations. Boxplots of absolute serum concentration of **(a)** arachidonic acid, **(b)** prostaglandin D2, and **(c)** prostaglandin E2 for all mucosal melanoma locations prior to any ICI therapy. Significant differences are indicated by an asterisk (*p* $\leq 0.05$). Trend differences are indicated by a cross (*p* $\leq 0.100$).

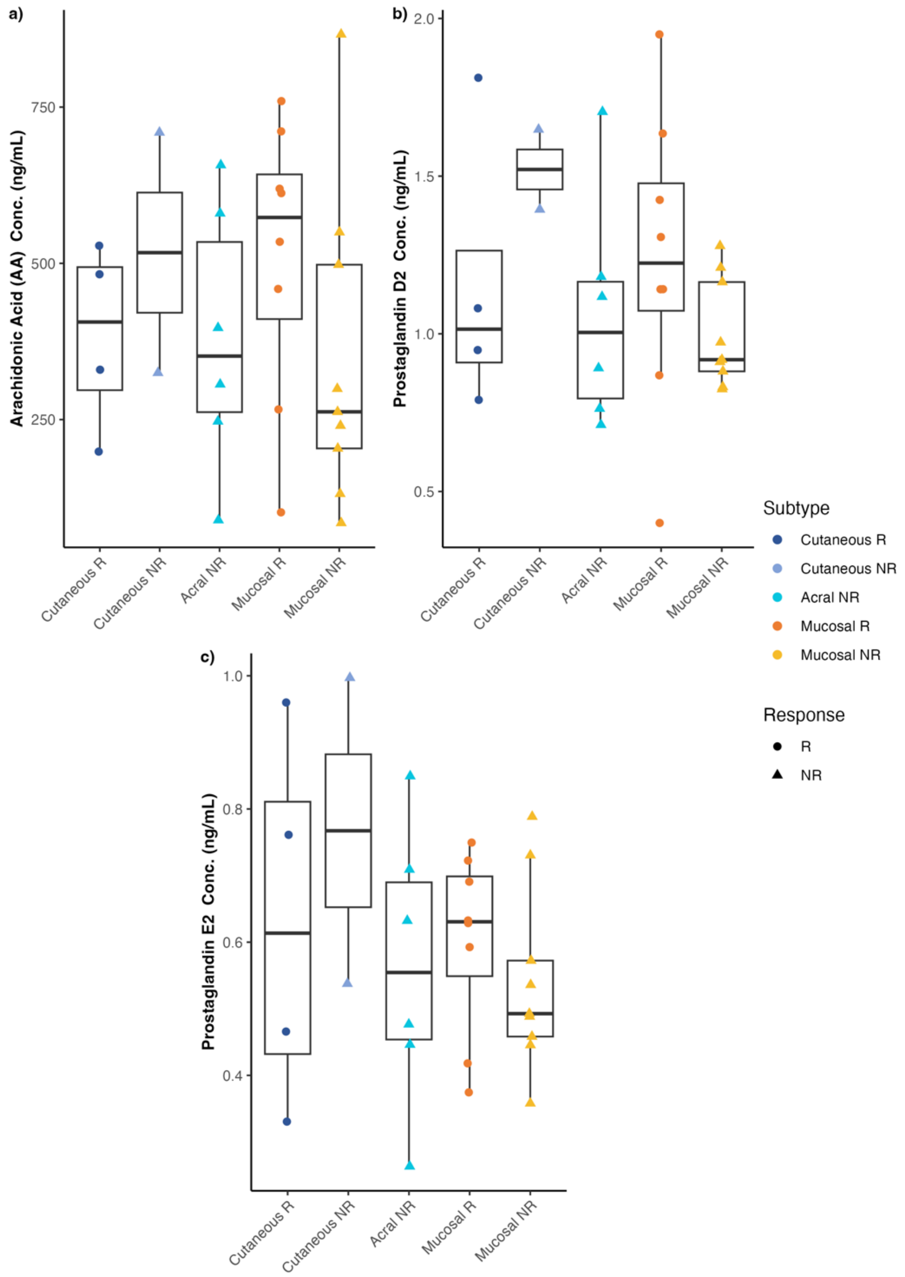


**Supplementary Figure 5.** Comparisons of major prostaglandins between ICI therapy responders and non-responders for each melanoma subtype. Boxplots of absolute serum concentration of **(a)** arachidonic acid, **(b)** prostaglandin D2, and **(c)** prostaglandin E2 for ICI therapy responders and non-responders for each melanoma subtype. Acral responders were not included in these plots as there were no acral responders in our study population. Significant differences are indicated by an asterisk ($p \leq 0.05$). Trend differences are indicated by a cross ($p \leq 0.100$).

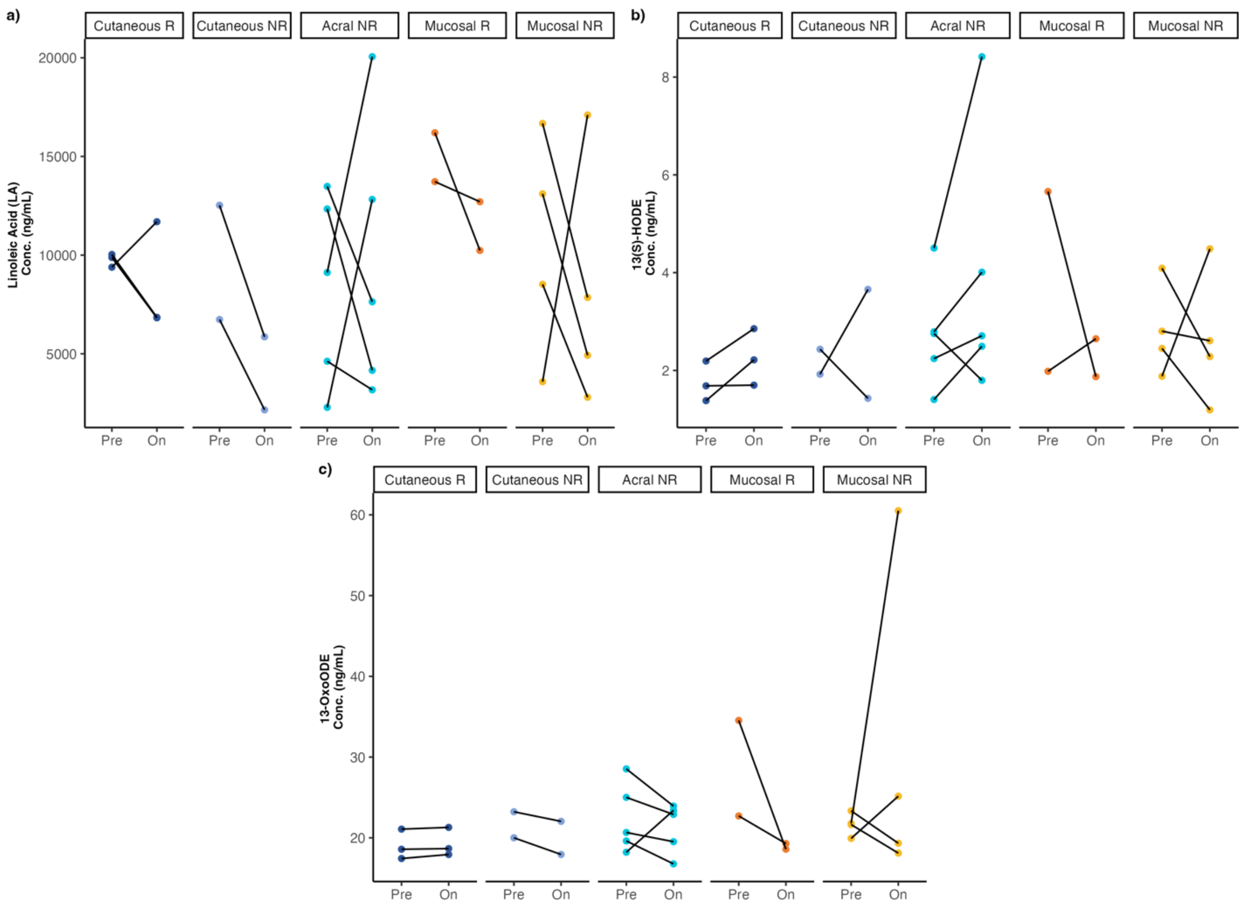


**Supplementary Figure 6.** Line graphs of linoleic acid-related metabolism oxylipins for matched pre- and on/post-treatment samples absolute serum concentration of **(a)** linoleic acid, **(b)** 13(S)-HODE, and **(c)** 13-OxoODE for ICI therapy responders and non-responders across melanoma subtypes prior to any ICI therapy.
